# Conformational footprinting of proteins using a combination of top-down electron transfer dissociation and ion mobility

**DOI:** 10.1101/283796

**Authors:** Albert Konijnenberg, Jinyu Li, Johny Habchi, Marion Dosnon, Giulia Rossetti, Rita Grandori, Sonia Longhi, Paolo Carloni, Frank Sobott

## Abstract

In recent years native mass spectrometry has been increasingly employed to study protein structure. As such a thorough understanding of the effect of the gas-phase on protein structure is becoming increasingly important. We show how a combination of top-down ETD and ion mobility can be used to probe the gas-phase structure of heterogeneous protein ensembles. By applying collisional activation to the non-covalently bound ETD products after IM separation, the peptide fragments can be released while maintaining the conformational information of the protein ion. We studied the unknown gas-phase structures of the measles virus (MeV) phosphoprotein X domain (P_XD_), which shows a wide range of different conformations in the gas-phase. We then generated structural models by state-of-the-art gas-phase steered molecular dynamics, which we verified using restraints from ion mobility and the fragment patterns observed. Our findings illustrate the applicability of ETD for obtaining conformational specific structural information on heterogeneous protein ensembles.

## Introduction

In recent years native mass spectrometry is increasingly used to study proteins and protein complexes. Although traditionally largely employed as a supportive technique, more recently native MS has shown to be able to address structural questions that are difficult to tackle with traditional biophysical techniques^1–4^. Such studies are based on the notion that proteins can be transferred natively into the gas-phase. Although for globular proteins a large body of evidence suggest that they can be transferred into the gas-phase whilst retaining their solution fold^5,6^, such conclusions might be harder to justify for proteins that have little secondary or tertiary structure. As such a thorough understanding of the effect of the gas-phase on protein structure is becoming increasingly important, but especially for structurally heterogeneous protein ensembles still remains difficult to obtain.

Two popular approaches to study protein structures in the gas-phase are ion-mobility mass spectrometry (IM-MS) and top-down radical-driven fragmentation experiments. Ion mobility adds the possibility to a mass spectrometry experiments to detect and separate different conformations and provide low resolution global structural information in the form of a collision cross section (CCS). These CCSs can then be compared to those obtained after calculations on structures obtained by X-ray crystallography or NMR spectroscopy^6^. Top-down experiments, on the other hand, are based on sequencing of intact proteins with non ergodic, radical-driven fragmentation methods combined with mass spectrometry detection, yielding information on the primary structure of a protein with amino acid resolution. Several studies have indicated that analysis of intact proteins under native conditions can also yield structural information, for example on the backbone flexibility or the exposed surface area (solvent accessibility of residues)^7–10^. For example, it was shown that top-down electron transfer dissociation (ETD) enables surface mapping of the tetrameric protein alcohol dehydrogenase, by combining information from product ion spectra with solvent accessibility prediction of the amino acids in the protein structure^10^. Furthermore combining field asymmetric ion mobility spectrometry (FAIMS) separation with ECD revealed that the fragmentation patterns observed can be linked to the solution structure of a protein^11^. Here we provide evidence that top-down ETD can also be used to simultaneously characterize conformational differences based on their mobility arrival time-dependent ETD product ion spectra, when used in conjunction with ion mobility.

When ETD is coupled with IM (Fig S1), the data can be viewed as a 2D intensity heat map, i.e. mobility arrival time versus mass-to-charge ratio (m/z). Specific regions of interest can be selected using procedures within the software to extract ETD product ion spectra from the heat map (Fig S1B). Product ions originating from ETD, but released before the ion mobility cell, can be easily selected in this way, as they tend to lie on a diagonal line of increasing arrival time and m/z in the 2D intensity heat map. Alternatively, the ETD product ions may be liberated after ion mobility (following supplemental activation) when non-covalent interactions keep them together^10,12^. Since the supplemental activation is applied upon entry into the transfer cell, positioned after ion mobility separation, fragments that are released at this stage will retain the mobility arrival time of their precursor (Fig S1B). This allows assignment of product ions to a specific precursor conformation, based on their common mobility arrival time. Thus, combining ion mobility with ETD allows for differentiation of fragments from co-existing conformers within a single experiment. Although (excessive) supplemental activation might cause conventional collision induced dissociation (CID), we observe that CID products (*b* and *y-*ions) are only a minority of the fragments observed, with the majority being *c* and *z* ions which can be easily distinguished from CID products based on their mass differences.

## Results

To test the ability of native top-down ETD to probe structural differences we turned to the well-characterized protein Calmodulin. In absence of Ca^2+^ Calmodulin has been shown to exist in equilibrium between a compact (globular) and an extended (“dumbbell”) conformation^13^ (Fig. 1A). Our experiments yielded virtually similar results to previous IM-MS experiments on Calmodulin in the gas phase in terms of absolute collision cross sections and intensities of the conformations^14^, proving that the presence of radical anions does not interfere with ion mobility experiments. Using this approach we applied ETD fragmentation to the 7+ and 8+ charge states of CaM, which both display the two main CaM conformations. Figure 1 shows the fragments observed for the 8+ charge state extracted from the arrival times of the compact conformation (Fig 1B) and extended conformation (Fig 1C) upon applying 40V of collisional activation after ETD/IM. Applying supplemental energy is crucial to release the fragments from the ETnoD products, most likely as non-covalent interactions are keeping the fragments in place^10,12^ A distinct difference in both ETD fragmentation efficiency and fragment pattern is observed between the fragments originating from the compact and the extended precursor (Fig. 1A). A similar trend is observed for the fragment patterns from both conformations in the 7+ charge state. Assignment of the observed fragments from the compact and extended conformations provided two different product ion spectra. The compact conformation yields fragments that originate mainly from the termini of the protein, whereas fragments from the extended conformation essentially cover the complete protein sequence (Fig. 1). Mapping the observed fragments onto the known three-dimensional structures of calmodulin shows for the compact conformation that the fragments mainly stem from loops and helices that are located on the exposed surface of the protein. For the extended conformation of CaM we observe the fragments to be much more widespread over the protein structure and even located on the extended linker helix (α4-α5), which is buried in the compact structure. As such we show that the main structural differences between these structures are reflected in the respective fragmentation patterns, validating the conformational specificity of native top-down ETD fragment patterns.

**Figure 1.**
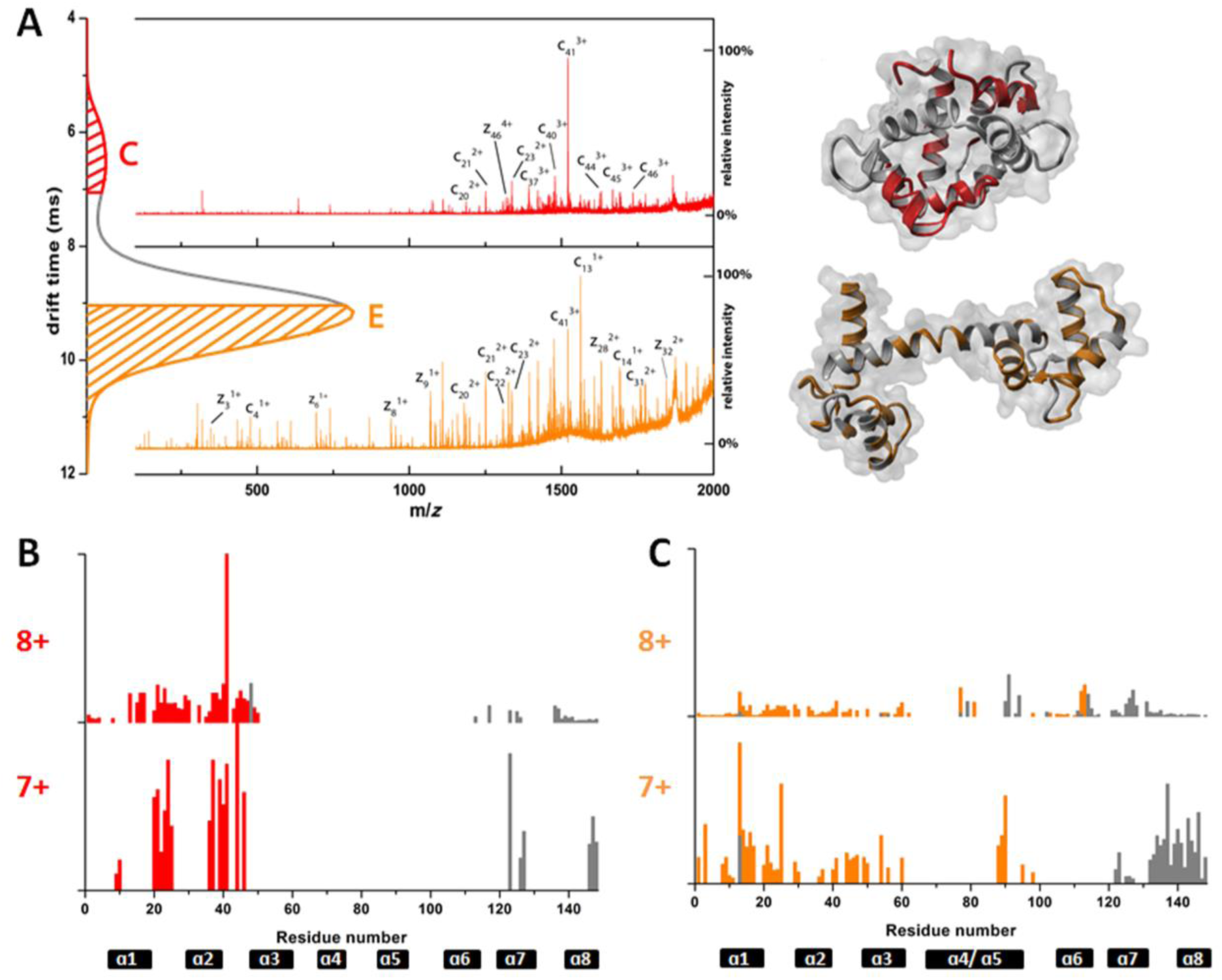
Conformational footprinting of the two conformations of Calmodulin. **A** conformational specific fragmentation spectra of each of the two conformations of Calmodulin, with their respective structures depicted on the right side. Fragments are shown on the structure of the structure in their respective colors. Fragmentation patterns for the 8+ and 7+ of **B** the compact, globular conformation and **C** the extended, dumbbell structure of CaM. Black bars indicate the location of the 8 alpha-helices that make up the structure of CaM.

Having showed that this method is able to yield different fragmentation patterns for different conformations, we aimed to apply it to a system where the gas-phase structure of the protein was unknown. Here we used native MS and top-down ETD to probe the unknown gas-phase structure of the three-helix bundle phosphoprotein X domain (P_XD_) from measles virus (MeV) in the gas phase. P_XD_ is required for MeV replication, via binding with the nucleoprotein C-terminal domain (N_TAIL_)^15^. P_XD_ has been studied by X-ray crystallography and NMR, yielding a compact three α-helix bundle structure^16,17^. In native mass spectrometry experiments P_XD_ displays a broad charge state distribution indicative of a protein that is structurally heterogeneous^18^, which is in line with recent work on the conformational landscape of P_XD_^19^. Ion mobility experiments showed that the 3 lowest charge states (4^+^-6^+^) all displayed multiple conformations (Fig. S2). Especially the 5^+^ charge state, with 5 distinct conformations, displays an astonishing degree of plasticity for a protein of only 55 amino acids (AA) (Fig. 2). Interestingly, these conformations appear to be stable in the gas phase over a 60 V range of ion acceleration voltages, in a collision-induced unfolding (CIU) study (Fig. S3).

**Figure 2.**
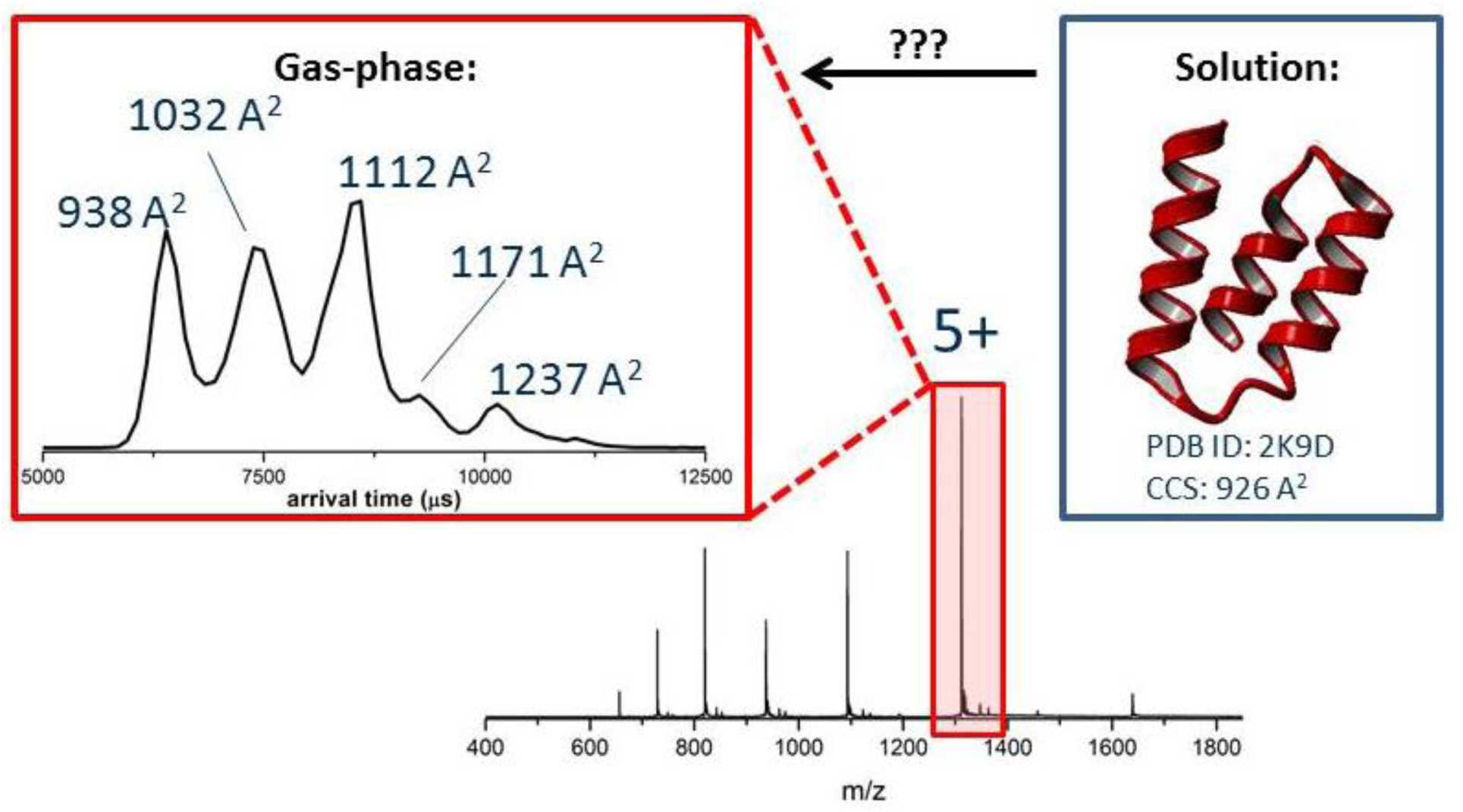
Mass spectra of P_XD_ reveal a broad charge state distribution indicative for an unfolded protein, as is reinforced by the multiple conformations observed for the 5^+^ charge state of P_XD_ (red inset). The blue inset shows the solution structure of P_XD_ as solved by NMR spectroscopy^20^, revealing it to be a stable three-helix bundle in solution.

We investigated the gas-phase structure of P_XD_ using native top-down ETD on the 5^+^-7^+^ charge states of P_XD_, combined with ion mobility to separate the different conformations. In order to release ETD fragments from the 5^+^ charge state, we required supplemental activation. The need for supplemental activation indicates that the higher-order structure is at least partially maintained in the gas phase, thus preventing the ETD fragments from being released. The requirement for supplemental activation was not observed for higher charge states. For both 6^+^ and 7^+^, we observed fragments without supplemental activation. This suggests that the more unfolded states of the proteins, which are observed at higher charge states, undergo a loss of secondary and tertiary structure. Further evidence for structural heterogeneity is in the broad charge state series observed for P_XD_ (4^+^-9^+^). The fragment patterns obtained for 6^+^ and 7^+^ charge states show fragments between from amino acids 1-21 and 40-55, with the 7^+^ charge state additionally showing fragments from amino acids 30-40 (Fig. S4). This is remarkable as these fragments stem from parts of the protein that appear as α-helixes in the solution structure^16^; α-helix 1 and α-helix 3 encompasses amino acids 4-12 and 33-47, respectively. The suggested loss of structure for higher charges states of P_XD_ is supported by the ion mobility data which yields collision cross section (CCS) between 1169 and 1334 Å^2^ for the 6^+^ and 1354 Å^2^ for the 7^+^ charge state (Fig. S2). Comparing these CCSs to the theoretical value of 960 Å^2^ for the solution structure indeed points towards a less compact, more open conformation of P_XD_ for those charge states.

We then focused on the 5^+^ charge state, which displayed a broad range of distinct conformations. We observed 5 conformers with CCSs of 938, 1032, 1112, 1171 and 1237 Å^2^, which are well separated by IM-MS (Fig. 2). By applying supplemental activation and correlating the product ions with the individual conformers via the mobility arrival time, we generated conformation-specific ETD fragment patterns. We ran a series of increasing supplemental activation (Fig.S5) to assess its effect on the release of fragments from P_XD_. At least 20V of supplemental activation was required in order to observe fragments. Increasing the supplemental activation further yielded a more extensive fragmentation pattern, although supplemental activation above 50V dramatically deteriorated the spectral quality. We observed extensive sequence coverage for the two more extended conformations (Fig. 3). Interestingly, similar to the 6^+^ and 7^+^ we did not obtain fragments between AA 23-29 for any of the conformations, which are part of α-helix 2 in the solution structure (Fig. 3). The difficulty to sequence α-helices by top-down ETD has been reported before, with specific salt-bridges - on top of the already extensive hydrogen bonding between residues in the helix - apparently strengthening the α-helix to an extent that it resists fragmentation even with supplemental activation^20^. For the third conformation we observed similar fragmentation patterns as for the two most extended conformations, with the exception of a large extent of α-helix 3, whereas the fragments continue to be observed from the loop between α-helix 2 and 3. For the two most compact structures we do not observe extensive fragmentation, but rather only some ETD fragments from the termini. Comparing these fragmentation patterns to the observed compact CCSs that these two species display, it is expected that significant higher-order structure is maintained for these conformations. As all conformations are activated under similar conditions, the differences in fragmentation patterns might point towards a loss of non-covalent interactions between or within the three-helix bundle in the gas-phase for the three most extended conformations, whereas the more compact conformations seem to retain enough interactions to keep the fragments together, even with supplemental activation applied. It should be noted that the P_XD_ conformations of the 5^+^ charge state are stable up to 60 V collision energy, whereas a maximum of 50 V was used to release the fragments.

**Figure 3.**
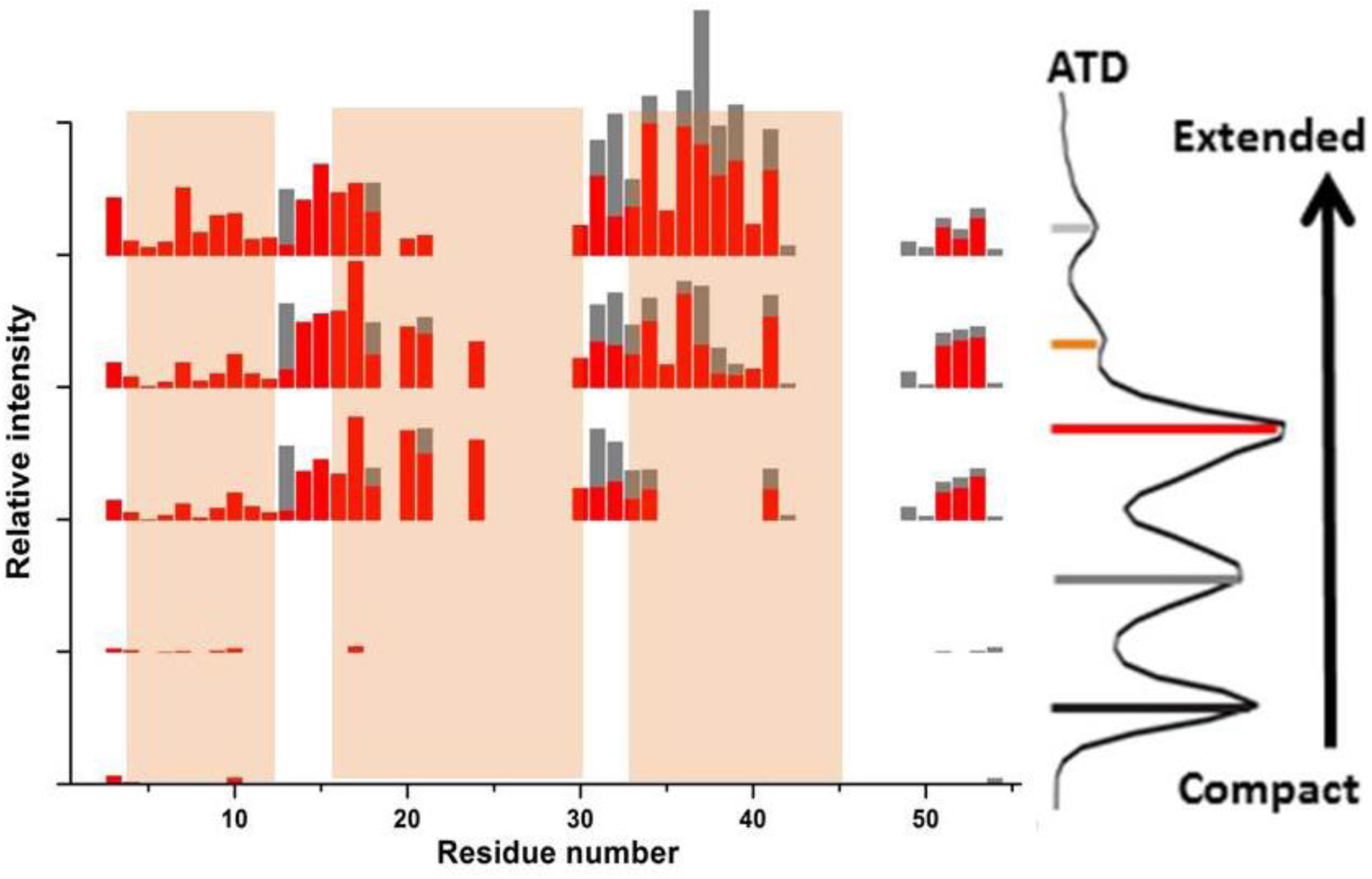
ETD fragments observed for the different conformations of the 5^+^ charge state of P_XD_. The mobility arrival time distribution (ATD) on the right shows the different conformations obtained from which the individual fragment patterns were extracted. C-fragments are colored red, with z-fragments in grey. Highlighted orange areas indicate alpha-helices in the solution structure of P_XD_.

To analyze the broad range of P_XD_ structures which we observed in the gas phase, we turned to computational modeling. First, we predicted the most probable protonation state of the protein with net charge state (5^+^) in the gas phase ([P_XD_]^5+^, Fig. S6). To this aim, we used a hybrid Monte Carlo/molecular dynamics (MC/MD) in-house protocol. The latter reproduces the electrospray ionization (ESI)-MS derived main charge state for folded proteins in the free state and in complex with other proteins^21,22^. Our calculations are based on a structural model of P_XD_ in solution (see Fig. S7), predicted from the NMR structure of the protein^23^. The resulting model is in good agreement with experimentally measured backbone N chemical shifts^16^ (Fig. S7C). Next, we used steered MD (SMD) simulations to investigate the partial unfolding of [P_XD_]^5+^ in the gas-phase. SMD has been successfully used to study the unfolding of a variety of proteins in solution^24^. In our calculations, the gyration radius of the protein (see Fig. S8A) - and hence the CCS that correlates with it (Fig. S8B) - was mildly steered during the dynamics. The calculated CCS features several plateaus, corresponding to a folded state (different from the native state, though) along with four partially unfolded states (Fig. 4A and Table S1). Within the approximate error margin - estimated to be 2-3%^25,26^, our calculated CCS values correlate well with the experimentally derived CCS, except for the value of the folded state, which is underestimated by 11% (Fig. 4A). Interestingly the calculated CCS for the folded protein corresponds closely to the most compact experimental CCS for the 4^+^ charge state of P_XD_, which is smaller than the theoretical CCS of the solution structure of P_XD_ (926 Å^2^), thus suggesting a compaction of the structure upon entering the gas phase. The [P_XD_]^5+^ conformations associated with the calculated CCS values (**I**-**V**) are displayed in Fig. 4. Because of the match between calculated (stable plateaus, i.e. regions where the radius of gyration remained constant) and experimental CCS, visual inspection of these conformations may give insights into the partial unfolding of the protein in the ESI-MS experiment and explain its observed ETD fragmentation patterns (Fig. 3). In the folded state, the three α-helix bundle slightly opens up (**I** in Fig. 4). This leads to the ejection of α-helix 3 (**II**). For both conformations **I** and **II**, which are rather compact, we do not see much ETD fragmentation. This is consistent with the conformers’ degree of secondary and tertiary structures. In **III**, α-helix 1 and α-helix 3 unravel. The latter is partially replaced by a new α-helix, closer to the C-terminus than α-helix 3. This replacement of helix 3 by one located closer to the C-terminus is consistent with the absence of fragments observed in the ETD experiments (Fig. 3 and 4B): strong interactions are maintained in this region, preventing a release of the fragments. In **IV** and **V**, α-helix 2 progressively unravels. A “silent” area, which is not sequenced, is observed between amino acids 22 and 28 throughout all conformations and levels of applied supplemental activation (Fig S5). Although α-helix 2 slowly starts to unravel for the two most extended conformations, this region maintains a strong α-helical character (Fig. 4B), possibly preventing the release of any fragments. A similar effect is observed for the area encompassing the newly formed α-helix near the C-terminus. The formation of this new α-helix is interesting, as it is in this region that poor sequence coverage is obtained for the three largest conformations. This “silent” area, which is not sequenced, is followed by an unstructured region that is fully sequenced. This is again in agreement with the fragmentation patterns observed for conformations **IV** and **V** (Fig. 4). These results validate that native top-down ETD combined with IM-MS provides significant structural information: the models for conformations **I-V** show to be consistent with experimental CCS data and furthermore provide a structural explanation for the ETD fragmentation patterns measured here.

**Figure 4.**
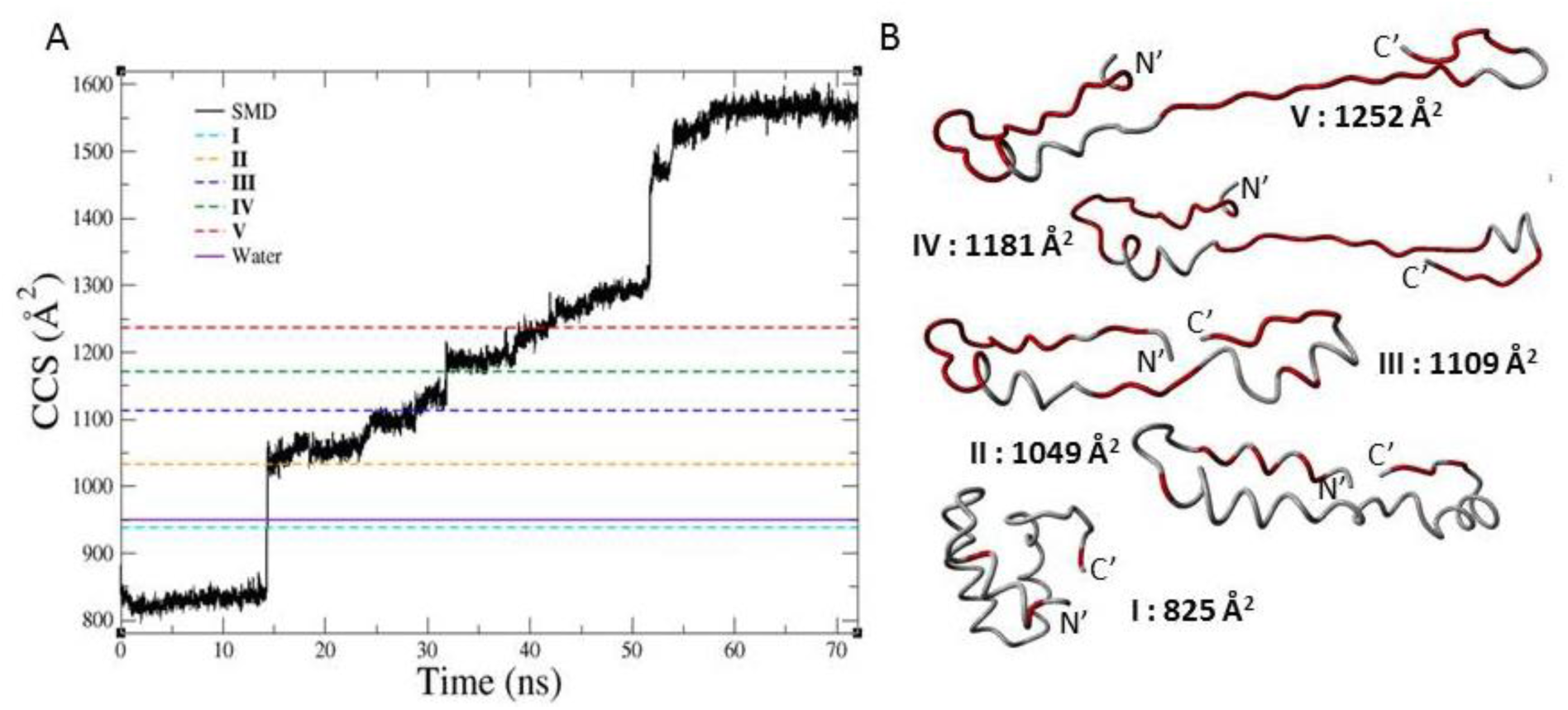
SMD simulations of [P_XD_]^5+^. **A** Calculated CCS values plotted as a function of simulated time. The experimental CCS values for conformations **I-V** are indicated by dashed lines and the calculated CCS value of the protein in aqueous solution is indicated by the solid line. **B** Most representative structural models of the different conformations of the 5^+^ charge states of P_XD_ based on the steered molecular dynamics simulations after cluster analysis of the structures present in the plateau areas, with the observed ETD fragments observed mapped in red for each ion-mobility separated conformation.

The observation that even at 50V of supplemental activation after ETD, we do not observe fragments from certain regions is interesting. Ever since the first research using radical driven fragmentation was applied to proteins, it was observed that supplemental activation is required to release fragments from the remaining complex^27–29^. However, as we go up stepwise in supplemental activation, we observe that after 30V the fragment pattern observed does not significantly change. This suggests that certain regions are simply not sequenced, or so stable that CID fragmentation occurs prior to release of these ETD fragments. As our results suggests that these regions maintain a high degree of secondary structure, a more systematic study on a well characterized protein structure should be performed to study the relation between gas-phase structure and the hierarchy of release of fragments with increasing supplemental activation.

To further validate our hypothesis that the behavior of P_XD_ is caused by unraveling of the three-helix bundle, we added the protein N_TAIL_ to P_XD_. The protein N_TAIL_ is a binding partner of P_XD_ and forms a 1:1 complex. Binding of the intrinsically disordered protein N_TAIL_ to P_XD_ is mediated by a disorder-to-order transition in the αMORE (molecular recognition elements) region of N_TAIL_, forming an α-helix that binds to the three-helix bundle motif of P_XD_ with an affinity in the micro molar range^30^. The interactions between P_XD_ and N_TAIL_ have been studied by IM-MS before, showing the formation of a 1:1 complex in the gas-phase^18^. As the three-helix bundle is required for binding of P_XD_ and N_TAIL_, we performed native top-down ETD on the N_TAIL_-P_XD_ complex, to see if a difference in the fragmentation pattern for P_XD_ can be observed. The fragile nature of the N_TAIL_-P_XD_ interaction is reflected in the dissociation of the complex in the gas-phase upon supplemental activation after ETD (Fig. S9A). However, as the ETD reaction takes place before the transfer cell, where dissociation can be induced, fragments observed should still reflect the nature of the N_TAIL_-P_XD_ complex. Top-down ETD of the native N_TAIL_-P_XD_ complex shows exclusively fragmentation at the N- and C-termini of P_XD_ (Fig. 5) over a broad range of collisional activation. These results are interesting as the much larger N_TAIL_ protein show a more extensive fragmentation, suggesting that the absence of fragments for P_XD_ is not caused by a low ETD fragmentation efficiency, but rather due to retention of the structure of P_XD_ (Fig. 5 and S9B). The fragment patterns of P_XD_ generated from its complex with N_TAIL_ resemble these observed for the two compact conformations of P_XD_ alone, where only fragments from the termini of P_XD_ were observed due to the retention of higher order structure. Similar as for PXD alone, the fragment pattern is not changed with increasing collision energy, further reinforcing a possible relationship between higher order structure and the observed ETD fragments.

**Figure 5.**
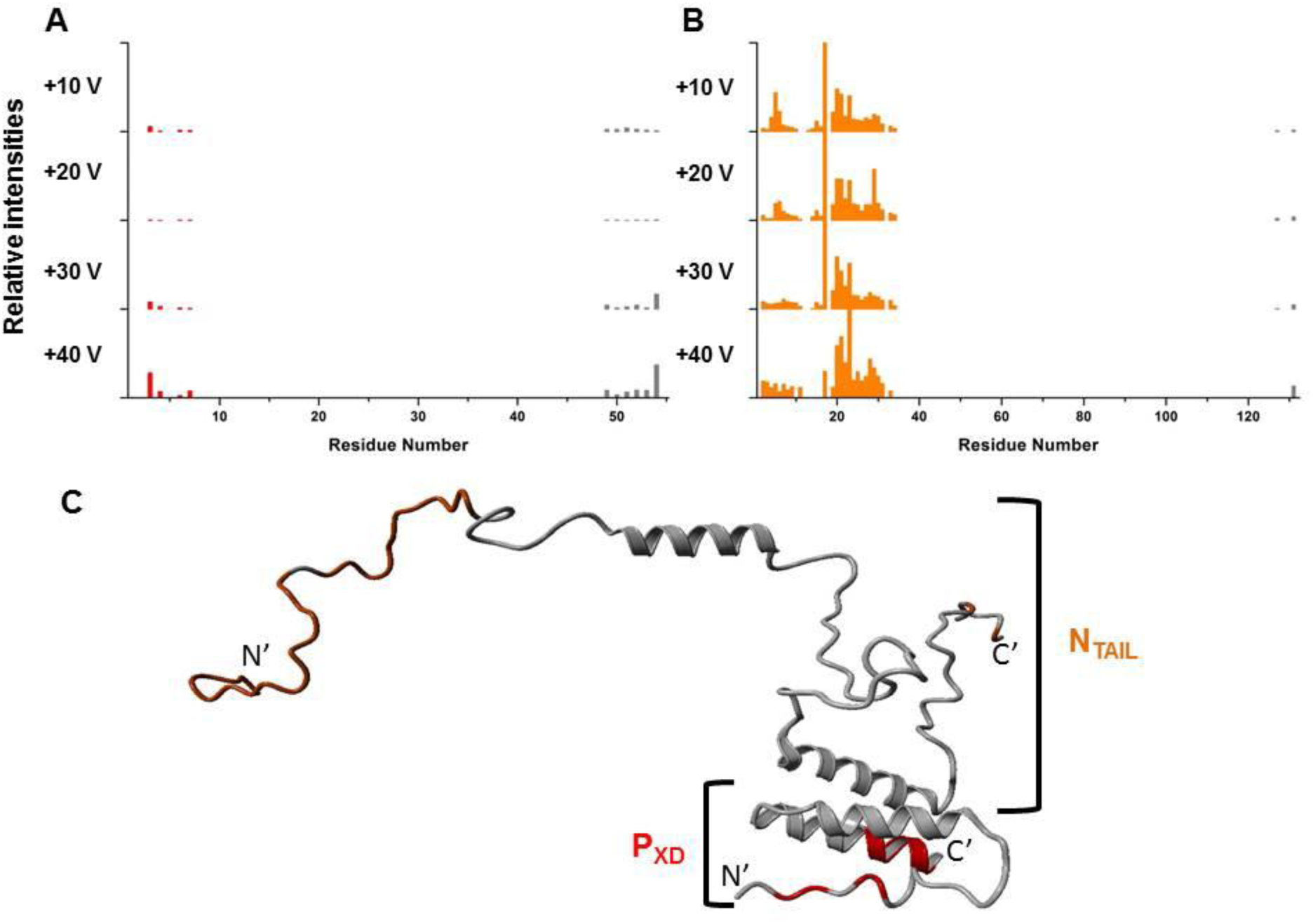
Fragment patterns for **A** P_XD_ and **B** N_TAIL_ generated by top-down ETD of the 10^+^ N_TAIL_-P_XD_ complex at various levels of supplemental activation reveal only terminal fragments, indicative of a stable three-helix bundle structure of P_XD_ within the complex. C-fragments are colored in red (P_XD_) or orange (N_TAIL_), whilst z-fragments are colored grey in both figures. **C** Solution model of the N_TAIL_-P_XD_ complex as calculated previously^18^ The structures of P_XD_ and N_TAIL_ are depicted in grey, with the observed ETD fragments mapped in red, whereas observed fragments for the intrinsically disordered protein N_TAIL_ are colored orange.

## Discussion

Here we present data in which ETD fragment ions have been generated from distinct conformational states of multiply charged protein ions within the same experiment. This method was employed to generate simplified spectra, each containing the product ions originating from the different more compact or elongated conformations of structurally heterogeneous proteins. Extraction of the relevant product ion spectra from the 2D heat map allows linking of sequence-specific information with individual conformations of the protein.

In conclusion we show that the top-down ETD fragmentation process is conformation-specific and can thus differentiate individually coexisting conformations of a protein in a single experiment. By doing so we established native top-down electron transfer dissociation as a powerful addition to the toolbox of mass spectrometry techniques for structural biology. Paired with ion mobility, native top-down fragmentation approaches can provide essential structural restraints, both globally at low resolution (distinguish conformations by CCS (Å^2^)) and per residue (determine solvent accessibility or flexibility). These restraints in turn could be used to refine or validate models generated from homology modeling or molecular dynamics^31,32^, or serve as selection criteria for possible candidates from large ensembles of structures, as can e.g. be generated with software such as the *Flexible Meccano* package for intrinsically disordered proteins^33^.

## Methods

### Sample preparation

The pDEST14 vector, allowing the expression of N-terminally hexahistidine tagged recombinant N_TAIL_ under the control of the T7 promoter, and the pDEST14 construct encoding C-terminally hexahistidine tagged MeV XD were used for the expression of the recombinant proteins. Expression and purification of the protein was carried out as described previously^16,23^, except that the final gel filtration step was carried out using 10 mM ammonium acetate pH 6.5 as elution buffer. To obtain the PXD-NTAIL complex an equimolar concentration of NTAIL was added, incubated for 5 minutes and subsequently diluted to a final concentration of 10 µmolar. Calmodulin from bovine testes (ID: P1431) was obtained from Sigma Aldrich (St Louis MO, US) and dissolved in 100mM Ammonium Acetate (AmAc) pH 6.8 to a concentration of 100uM. Sample was subsequently aliquoted and stored at −20C for further use. CaM aliquots were diluted to a final concentration of 10uM in 100mM AmAc and buffer exchanged twice on a Biorad (Hercules CA, US) P6 spin column (ID: 732-6221) against 100mM AmAc to obtain calcium free calmodulin sample. Final samples were measured immediately after preparation

### Mass spectrometry

Multiply charged ions were generated using nano-electrospray ionization with in-house prepared gold-coated capillaries and a capillary voltage of +1.5 kV. All experiments were performed on a Waters Synapt G2 HDMS with ETD option. Important voltages were: sampling cone 25-40 V, extraction cone 1 V, trap collision energy 4 V, transfer collision energy 70 V for CaM and 50 V for P_XD_ (unless indicated otherwise) and trap DC bias 30 V. Selected charge states for CaM, P_XD_ and N_TAIL_-P_XD_ were mass-selected using the 32 k quadrupole and underwent an ion-ion ETD reaction with reagent 1,4 dicyanobenzene in the trap T-wave (at ∼5×10^-2^ mbar He) of the instrument. These ETD ions then pass through the T-wave ion mobility stage (at ∼1.5 mbar N_2_) and are subjected to CID in the transfer T-wave (at ∼2×10^-2^ mbar Ar) and then analyzed using TOF-MS. Ion mobility separation was performed at a wave velocity of 300 m/s and a wave height of 13 V. ETD settings used were a scan time interval of 1 s and a reagent refill time of 0.1 s. The wave velocity in the trap cell, where the ETD fragmentation takes place, was 300 m/s with a wave height of 0.25 V. Data acquisition and processing were carried out using MassLynx4.1. Peak assignment was performed by manually comparing the observed fragment masses with an in-silico generated fragment list from the protein sequence. A cutoff of 3 times signal-to-noise or greater was used to determine which fragments were considered to be observed. Relative intensities of fragments were obtained by normalizing the fragment intensities against the most intense fragment observed. For spectra with multiple conformations another normalization round was performed, which scaled each intensity relative to the area under the drift time curve of the conformation it was found in.

Experimental collision cross sections (CCS) were determined by calibration against denatured proteins of known CCS (cytochrome C, ubiquitin and myoglobin) as reported elsewhere^34^. Denatured proteins were chosen as their charge and CCS increment reflects the unfolding of P_XD_ in the gas-phase and the intrinsically disordered nature of N_TAIL_ better than native globular proteins would.

### Computational modeling

Based on the solution NMR structure of P_XD_ (residues 462-505, PDB ID: 2K9D^23^), 200 initial models of P_XD_ (residues 459-507 with attached C-terminal hexa-histidine tag) were generated using MODELLER 9v9 package^35^. The lowest-DOPE (discrete optimized protein energy) score^36^ model was selected as the starting structure for the following MD-based refinement in solution. The protonation states of residues in solution were assigned according to the corresponding pK_a_ values calculated by using the H++ webserver^37^. P_XD_ was inserted into a water box with edges of 50×58×51 Å^3^ containing ∼4,420 water molecules and 5 Cl^-^ ions to neutralize the system. The AMBER ff99SB-ILDN force field^38–41^ and TIP3P force field^42^ were used for the protein and Cl^-^ ions, and for water, respectively. Periodic boundary conditions were applied. Electrostatic interactions were calculated using the Particle Mesh Ewald (PME) method^43^, and the cutoff for the real part of the PME and for the van der Waals interactions was set to 1.0 nm. All bond lengths were constrained using the LINCS algorithm^44^. Constant temperature and pressure conditions were achieved by coupling the systems with a Nosé-Hoover thermostat^45,46^ and an Andersen-Parrinello-Rahman barostat^47^. A time-step of 2 fs was employed. The protein underwent 1000 steps of steepest-descent energy minimization with 1000 kJ mol^-1^ nm^-2^ harmonic position restraints on the protein complex, followed by 2500 steps of steepest-descent and 2500 steps of conjugate-gradient minimization without restraints. The system was then gradually heated from 0 K up to 300 K in 20 steps of 2 ns. 200 ns long MD simulation at 300 K and 1 atm pressure was carried out using GROMACS 4.5.5^48^. The structure closest to the average conformation of the protein in aqueous MD simulation, which is in good agreement with the experimentally measured chemical shifts^16^ (Fig. S6), was employed as starting structure for the gas-phase simulations. The solvent and the Cl^-^ counterions molecules were removed.

A hybrid Monte Carlo (MC)/MD protocol^21,22^ was used to identify the most probable protonation states of P_XD_ at 5^+^ ([P_XD_]^5+^) in the gas phase. The protocol is based on OPLS/AA^49^ force field energy augmented by additional energy terms associated with the gas-phase basicity (GB) of ionizable residues^21^ (see ref.^22^ for detailed description). After the MC/MD calculation reached convergence (Fig. S5), the selected aqueous structure with the identified lowest energy protonation state for the [P_XD_]^5+^ underwent steered molecular dynamics (SMD) simulation at 300 K for 72 ns in the gas phase to enforce the protein to extended conformation by pulling the gyration radius (R_g_) of the protein (see Fig. S8). The SMD spring constant was set as 500 kJ•mol^-1^•nm^-2^, the velocity is 0.0001 nm per 1000 steps and the target R_g_ value is 4.5 nm. These values are similar to those applied in ref. ^50^. The SMD was carried out using GROMACS 4.5.5^48^ and PLUMED plug-in^50^ with the same setup as the one described for the aqueous MD simulation, except that the time step was 1.5 fs and the force field was OPLS/AA^49^. Theoretical CCS values were calculated for structures every 15 ps using the projection approximation (PA) method implemented in the MOBCAL code^51,52^. Since the PA CCS (CCS_PA_) typically underestimates experimental CCS measured in nitrogen buffer gas by ∼14%^53^, the scaled values (CCS_exp_=1.14×CCS_PA_) were used for comparison with experimental results.

## Acknowledgements

The authors would like to thank the Hercules foundation for funding the Synapt G2 instrument.

## Author contributions

A.K. and F.S conceptually conceived the research. J.H. and M.D. produced and purified the proteins. A.K. performed the mass spectrometry experiments. J.L. and G.R. performed the SMD calculations. All authors contributed to interpreting the data and writing the paper.

## REFERENCES

1. Uetrecht, C., Barbu, I. M., Shoemaker, G. K., van Duijn, E. & Heck, A. J. R. Interrogating viral capsid assembly with ion mobility-mass spectrometry. Nat. Chem. 3, 126–32 (2011).

2. Zhou, M. et al. Ion mobility-mass spectrometry of a rotary ATPase reveals AT-induced reduction in conformational flexibility. Nat. Chem. 6, 208–15 (2014).

3. Konijnenberg, A. et al. Global structural changes of an ion channel during its gating are followed by ion mobility mass spectrometry. Proc. Natl. Acad. Sci. 111, 17170–17175 (2014).

4. Stengel, F. et al. Dissecting Heterogeneous Molecular Chaperone Complexes Using a Mass Spectrum Deconvolution Approach. Chem. Biol. 19, 599–607 (2012).

5. Hall, Z. & Robinson, C. V. Do charge state signatures guarantee protein conformations? J. Am. Chem. Soc. 23, 1161–1168 (2012).

6. Jurneczko, E. & Barran, P. E. How useful is ion mobility mass spectrometry for structural biology? The relationship between protein crystal structures and their collision cross sections in the gas phase. Analyst 136, 20–8 (2011).

7. Zhang, H., Cui, W., Wen, J., Blankenship, R. E. & Gross, M. L. Native electrospray and electron-capture dissociation in FTICR mass spectrometry provide top-down sequencing of a protein component in an intact protein assembly. J. Am. Soc. Mass Spectrom. 21, 1966–8 (2010).

8. Zhang, H., Cui, W. & Gross, M. L. Native electrospray ionization and electron-capture dissociation for comparison of protein structure in solution and the gas phase. Int. J. Mass Spectrom. 354–355, (2013).

9. Li, H., Wolff, J. J., Van Orden, S. L. & Loo, J. A. Native top-down electrospray ionization-mass spectrometry of 158 kDa protein complex by high-resolution Fourier transform ion cyclotron resonance mass spectrometry. Anal. Chem. 86, 317–20 (2014).

10. Lermyte, F. et al. ETD allows for native surface mapping of a 150 kDa noncovalent complex on a commercial Q-TWIMS-TOF instrument. J. Am. Soc. Mass Spectrom. 25, 343–350 (2014).

11. Robinson, E. W., Leib, R. D. & Williams, E. R. The role of conformation on electron capture dissociation of ubiquitin. J. Am. Soc. Mass Spectrom. 17, 1469–79 (2006).

12. Zhang, Z., Browne, S. J. & Vachet, R. W. Exploring salt bridge structures of gas-phase protein ions using multiple stages of electron transfer and collision induced dissociation. J. Am. Soc. Mass Spectrom. 25, 604–13 (2014).

13. Zhang, M. & Yuan, T. Molecular mechanisms of calmodulin’s functional versatility. Biochem. Cell Biol. (2011).

14. Wyttenbach, T., Grabenauer, M., Thalassinos, K., Scrivens, J. H. & Bowers, M. T. The effect of calcium ions and peptide ligands on the relative stabilities of the calmodulin dumbbell and compact structures. J. Phys. Chem. B 114, 437–47 (2010).

15. Habchi, J. & Longhi, S. Structural disorder within paramyxovirus nucleoproteins and phosphoproteins. Mol. Biosyst. 8, 69–81 (2012).

16. Johansson, K. et al. Crystal structure of the measles virus phosphoprotein domain responsible for the induced folding of the C-terminal domain of the nucleoprotein. J. Biol. Chem. 278, 44567–73 (2003).

17. Gely, S. et al. Solution structure of the C-terminal X domain of the measles virus phosphoprotein and interaction with the intrinsically disordered C-terminal domain of the nucleoprotein. J. Mol. Recognit. 23, 435–47 (2010).

18. DUrzo, A. et al. Molecular Basis for Structural Heterogeneity of an Intrinsically Disordered Protein Bound to a Partner by Combined ESI-IM-MS and Modeling. J. Am. Soc. Mass Spectrom. (2014). doi:10.1007/s13361-014-1048-z

19. Bonetti, D. et al. Identification and Structural Characterization of an Intermediate in the Folding of the Measles Virus X domain. J. Biol. Chem. M116.721126- (2016). doi:10.1074/jbc.M116.721126

20. Breuker, K., Bruschweiler, S. & Tollinger, M. Electrostatic stabilization of a native protein structure in the gas phase. Angew. Chem. Int. Ed. Engl. 50, 873–7 (2011).

21. Marchese, R., Grandori, R., Carloni, P. & Raugei, S. A computational model for protein ionization by electrospray based on gas-phase basicity. J. Am. Soc. Mass Spectrom. 23, 1903–10 (2012).

22. Li, J. et al. Molecular simulation-based structural prediction of protein complexes in mass spectrometry: the human insulin dimer. PLoS Comput. Biol. 10, e1003838 (2014).

23. Bernard, C. et al. Interaction between the C-terminal domains of N and P proteins of measles virus investigated by NMR. FEBS Lett. 583, 1084–9 (2009).

24. Hsin, J., Strumpfer, J., Lee, E. H. & Schulten, K. Molecular origin of the hierarchical elasticity of titin: simulation, experiment, and theory. Annu. Rev. Biophys. 40, 187–203 (2011).

25. Politis, A. et al. A mass spectrometry-based hybrid method for structural modeling of protein complexes. Nat. Methods 11, 403–6 (2014).

26. Politis, A. et al. Integrating ion mobility mass spectrometry with molecular modelling to determine the architecture of multiprotein complexes. PLoS One 5, e12080 (2010).

27. Horn, D. M., Ge, Y. & McLafferty, F. W. Activated ion electron capture dissociation for mass spectral sequencing of larger (42 kDa) proteins. Anal. Chem. 72, 4778–84 (2000).

28. Zubarev, R. A. et al. Electron Capture Dissociation for Structural Characterization of Multiply Charged Protein Cations. Anal. Chem. 72, 563–573 (2000).

29. Zubarev, R. A. et al. Electron Capture Dissociation of Gaseous Multiply-Charged Proteins Is Favored at Disulfide Bonds and Other Sites of High Hydrogen Atom Affinity. J. Am. Chem. Soc. 121, 2857–2862 (1999).

30. Dosnon, M. et al. Demonstration of a Folding after Binding Mechanism in the Recognition between the Measles Virus N TAIL and X Domains. ACS Chem. Biol. 10, 795–802 (2015).

31. Webb, B. et al. Modeling of proteins and their assemblies with the Integrative Modeling Platform. Methods Mol. Biol. 1091, 277–95 (2014).

32. Politis, A., Park, A. Y., Hall, Z., Ruotolo, B. & Robinson, C. V. Integrative modelling coupled with ion mobility mass spectrometry reveals structural features of the clamp loader in complex with single stranded DNA binding protein. J. Mol. Biol. 425, 4790–4781 (2013).

33. Ozenne, V. et al. Flexible-meccano: a tool for the generation of explicit ensemble descriptions of intrinsically disordered proteins and their associated experimental observables. Bioinformatics 28, 1463–70 (2012).

34. Bush, M. F. et al. Collision cross sections of proteins and their complexes: A calibration framework and database for gas-phase structural biology. Anal. Chem. 82, 9557–9565 (2010).

35. Marti-Renom, M. A. et al. Comparative protein structure modeling of genes and genomes. Annu. Rev. Biophys. Biomol. Struct. 29, 291–325 (2000).

36. Shen, M.-Y. & Sali, A. Statistical potential for assessment and prediction of protein structures. Protein Sci. 15, 2507–24 (2006).

37. Gordon, J. C. et al. H++: a server for estimating pKas and adding missing hydrogens to macromolecules. Nucleic Acids Res. 33, W368–71 (2005).

38. Lindorff-Larsen, K. et al. Improved side-chain torsion potentials for the Amber ff99SB protein force field. Proteins 78, 1950–8 (2010).

39. Pavelites, J. J., Gao, J., Bash, P. A. & Mackerell, A. D. A molecular mechanics force field for NAD+ NADH, and the pyrophosphate groups of nucleotides. J. Comput. Chem. 18, 221–239 (1997).

40. Walker, R. C., de Souza, M. M., Mercer, I. P., Gould, I. R. & Klug, D. R. Large and Fast Relaxations inside a Protein: Calculation and Measurement of Reorganization Energies in Alcohol Dehydrogenase. J. Phys. Chem. B 106, 11658–11665 (2002).

41. Craft, J. W. & Legge, G. B. An AMBER/DYANA/MOLMOL phosphorylated amino acid library set and incorporation into NMR structure calculations. J. Biomol. NMR 33, 15–24 (2005).

42. Jorgensen, W. L., Chandrasekhar, J., Madura, J. D., Impey, R. W. & Klein, M. L. Comparison of simple potential functions for simulating liquid water. J. Chem. Phys. 79, 926 (1983).

43. Darden, T., York, D. & Pedersen, L. Particle mesh Ewald: An N-log(N) method for Ewald sums in large systems. J. Chem. Phys. 98, 10089 (1993).

44. Hess, B., Bekker, H., Berendsen, H. J. C. & Fraaije, J. G. E. M. LINCS: A linear constraint solver for molecular simulations. J. Comput. Chem. 18, 1463–1472 (1997).

45. Hoover, W. Canonical dynamics: Equilibrium phase-space distributions. Phys. Rev. A 31, 1695–1697 (1985).

46. Nose, S. A molecular dynamics method for simulations in the canonical ensemble. Mol. Phys. 52, 255–268 (2006).

47. Parrinello, M. Polymorphic transitions in single crystals: A new molecular dynamics method. J. Appl. Phys. 52, 7182 (1981).

48. Berendsen, H. J. C., van der Spoel, D. & van Drunen, R. GROMACS: A message-passing parallel molecular dynamics implementation. Comput. Phys. Commun. 91, 43–56 (1995).

49. Jorgensen, W. L. & Tirado-Rives, J. Potential energy functions for atomic-level simulations of water and organic and biomolecular systems. Proc. Natl. Acad. Sci. U. S. A. 102, 6665–70 (2005).

50. Bonomi, M. et al. PLUMED: A portable plugin for free-energy calculations with molecular dynamics. Comput. Phys. Commun. 180, 1961–1972 (2009).

51. Shvartsburg, A. A., Mashkevich, S. V., Baker, E. S. & Smith, R. D. Optimization of algorithms for ion mobility calculations. J. Phys. Chem. A 111, 2002–2010 (2007).

52. Mesleh, M. F., Hunter, J. M., Shvartsburg, A. A., Schatz, G. C. & Jarrold, M. F. Structural Information from Ion Mobility Measurements: Effects of the Long-Range Potential. J. Phys. Chem. 100, 16082–16086 (1996).

53. Hall, Z., Politis, A., Bush, M. F. & Robinson, C. V. Charge state dependent compaction and dissociation of protein complexes: insights from ion mobility and molecular dynamics. J. Am. Chem. Soc. 134, 3429–3438 (2012).

