## Supplementary Materials for "Conformational footprinting of proteins using a combination of top-down electron transfer dissociation and ion mobility"

### Supplemental information

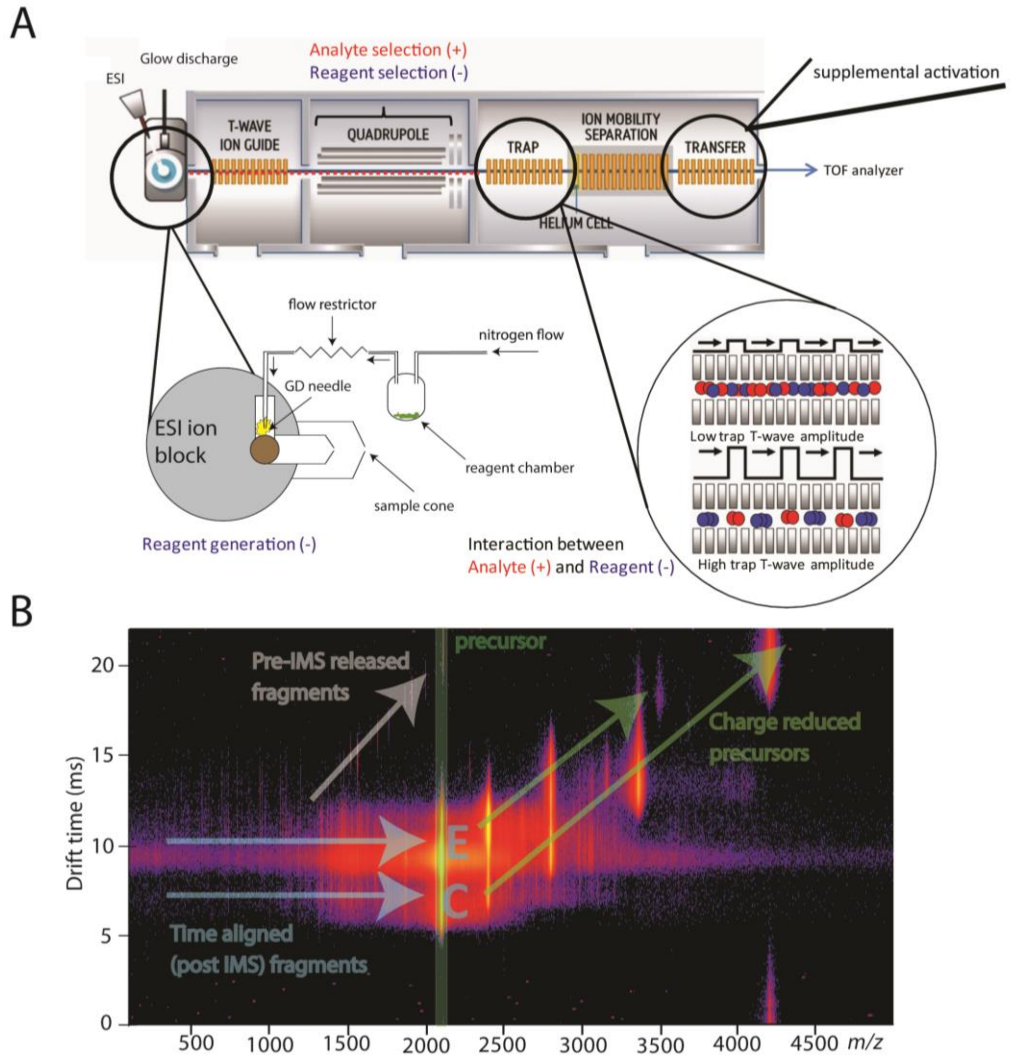


**Figure S1 A.** Schematic representation of the Synapt G2 instrument with ETD option. ETD reagent vapor is transferred to the low pressure region in the ion source block where glow discharge ionization generates radical anions. The ion mode polarity of the instrument is alternated as far as the trap cell to transfer both cationic analyte and anionic radical ions to the trap cell. In the trap cell, both populations of ions are allowed to mix and react to form both ETD and charge reduced products. Mixing is governed by the wave height. Low wave heights allow ions to roll over the 'waves' and mix, whereas high wave heights keep the ion separated and thus prevent reaction. **B.** Ion mobilogram of native top-down ETD-IM-MS of the 8+ charge state of CaM. Combining ETD with ion mobility allows for easy assignment of time-aligned fragments. When fragments are formed but not released from the conformational precursor yet, post-IMS supplemental activation can release them. As a consequence, these fragments (grey arrows, 'time aligned fragments'), will maintain the arrival time of their precursor conformation thus allowing conformational footprinting by ETD.

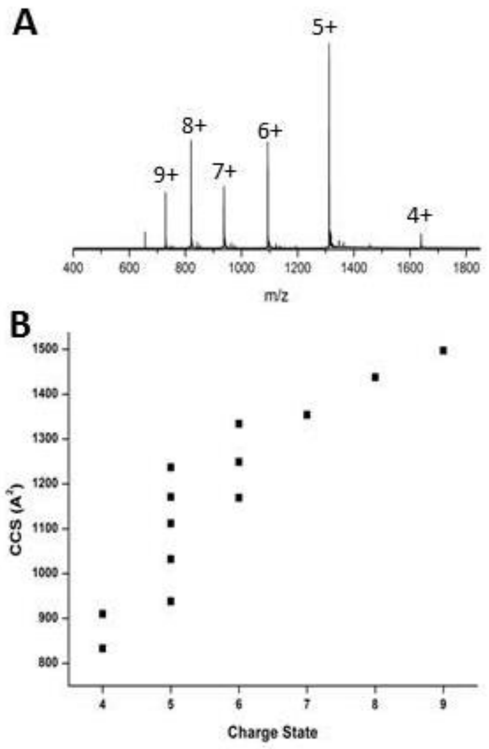


**Figure S2** $P_{XD}$ displays a broad range of conformations under native conditions (10mM Ammonium Acetate of pH 6.5). **A** Nano-ESI-IM-MS under non-denaturing conditions shows a broad charge state distribution, which is characteristic for proteins that are structurally heterogeneous. **B** IM-MS shows that $P_{XD}$ adopts a wide range of conformations, ranging from compact structures for the lower charge states, to elongated structures for the higher charge states which are typical for an unfolded protein.

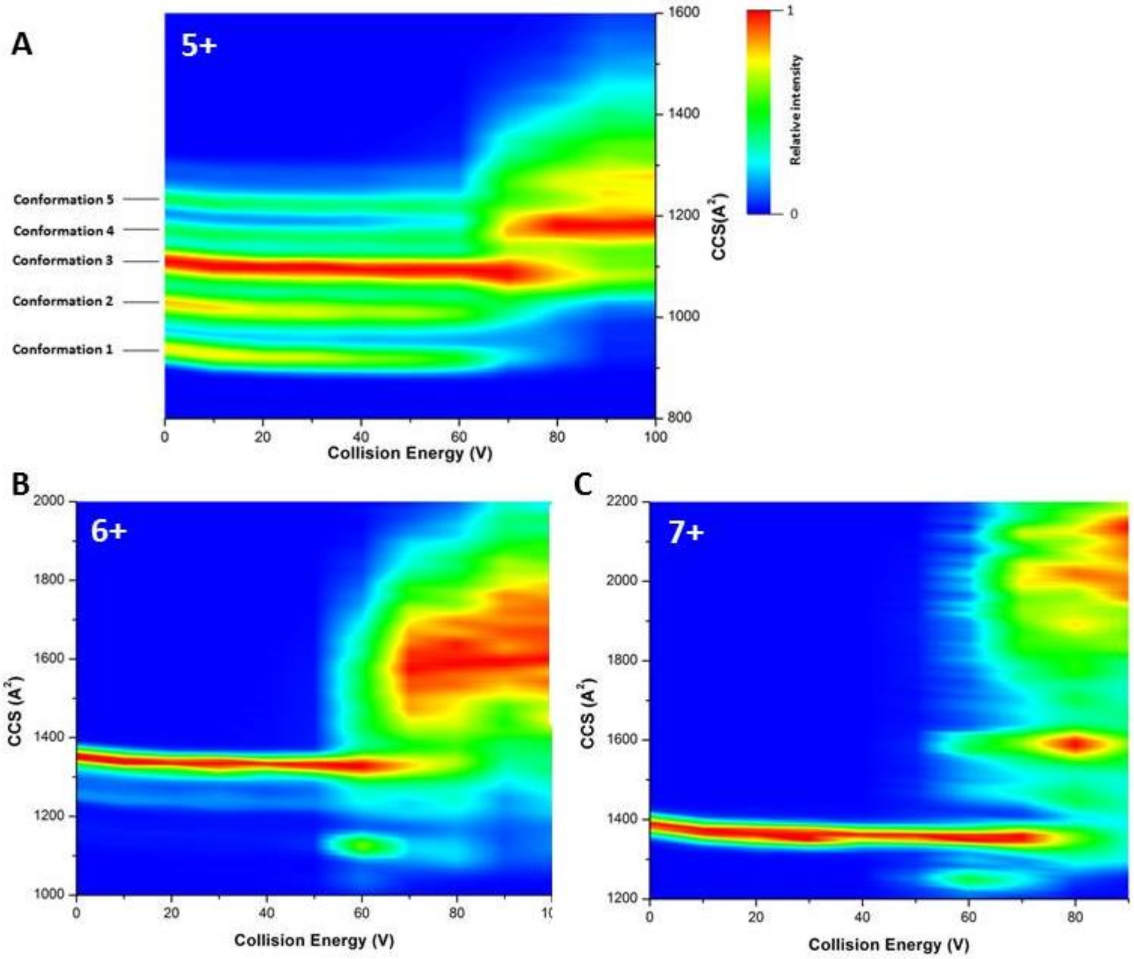


**Figure S3** Heat map of the collision induced unfolding (CIU) of the **A** $5^+$ charge state of $P_{XD}$. The 5 conformations observed are stable over a range of 60 V of ion acceleration energy (equivalent to 300 eV collisional activation), before displaying CIU. Similar unfolding trends are observed for the **B** 6+ and **C** 7+ charge state. Collision energies higher than 50V lead to a deterioration of spectral quality, most likely due to fragmentation of the protein by CID.

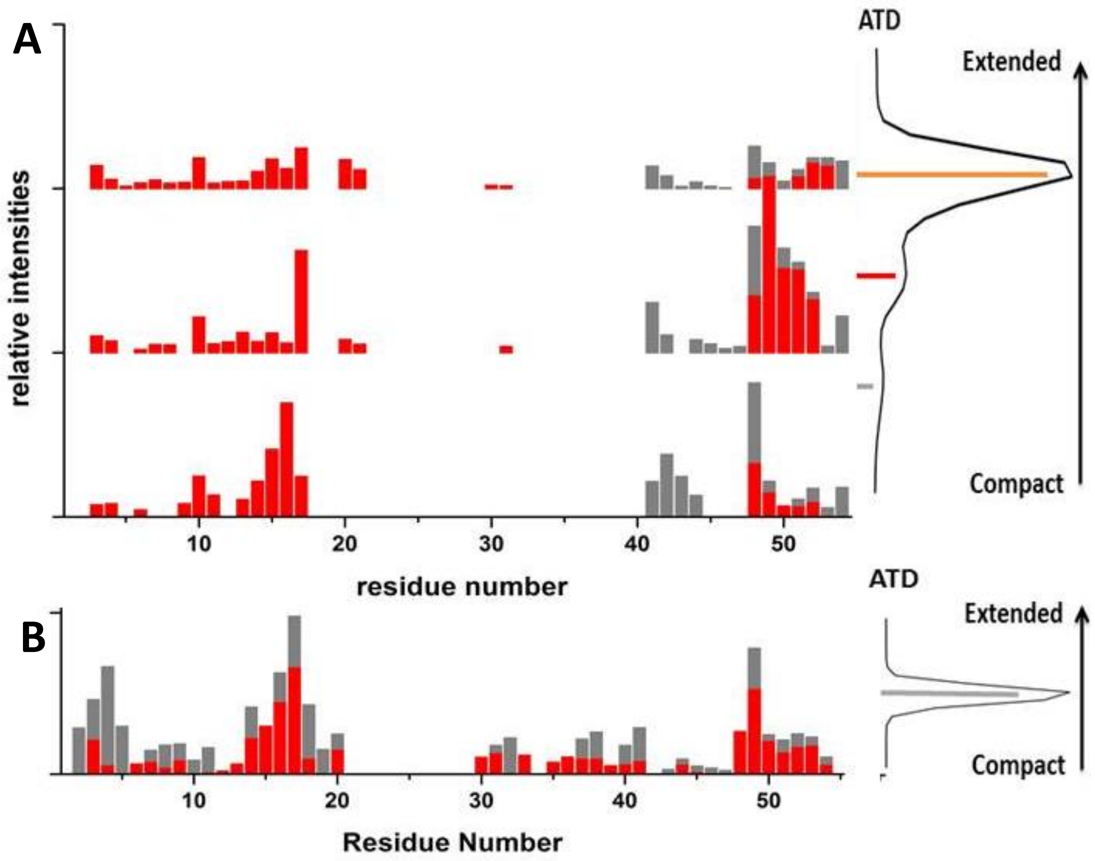


**Figure S4** ETD fragment patterns for the $6^+$ and $7^+$ charge state of $P_{XD}$. **A** ETD fragments for the 3 different conformations of the $6^+$ charge state of $P_{XD}$ obtained when 40 V supplemental activation is applied. **B** ETD fragment pattern for the $7^+$ charge state, which has a single conformation, under the same conditions as for the $6^+$ charge state.

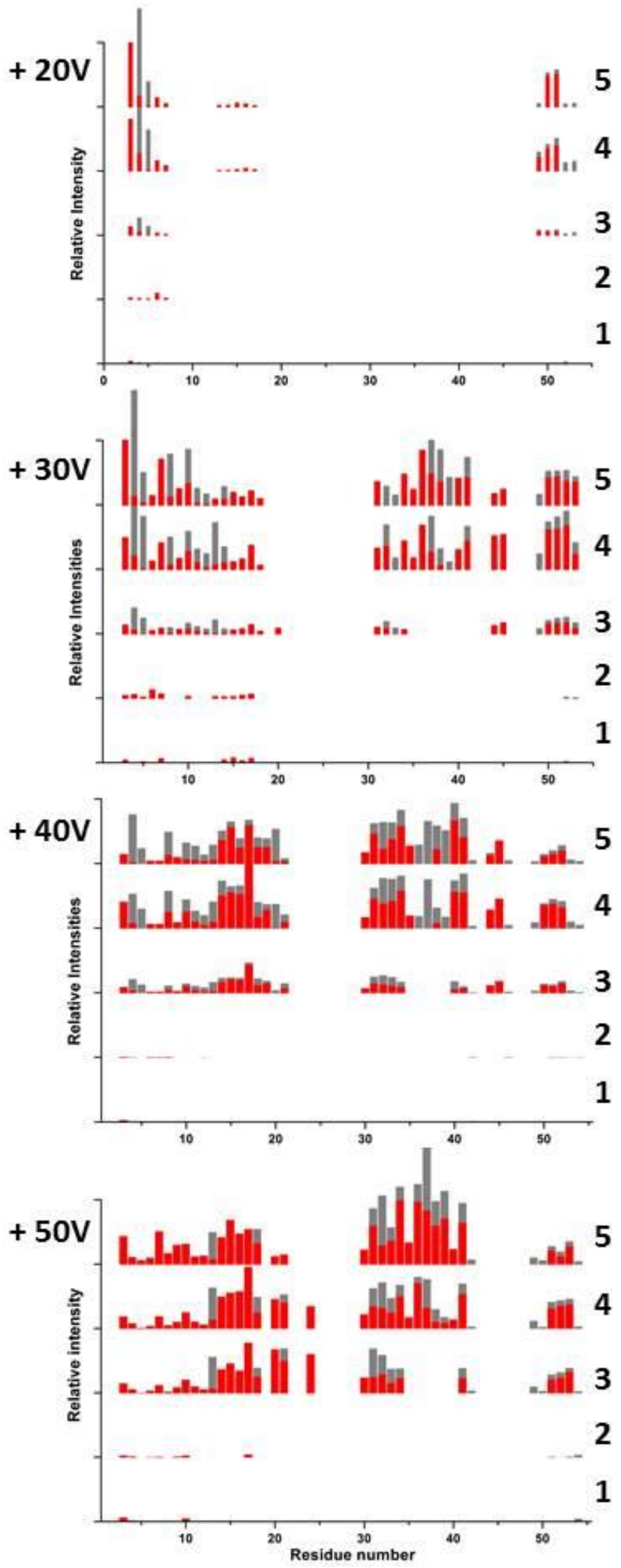


**Figure S5** Fragment patterns for each of the 5 conformations of the 5+ charge state of $P_{XD}$ (with 1 being the most compact and 5 the most extended conformation) with increasing supplemental activation.

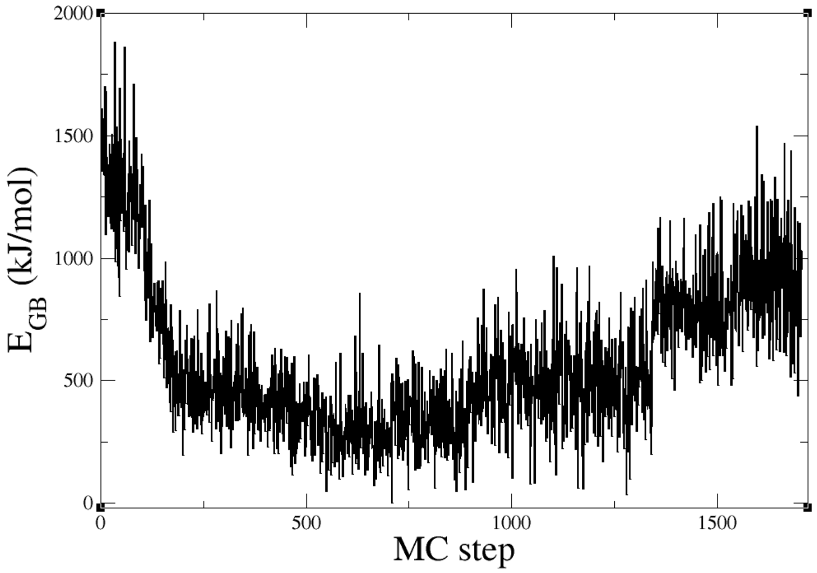


**Figure S6** Gas-phase basicity corrected energy ($E_{GB}$) of $[P_{XD}]^{5+}$ plotted as a function of MC steps in our combined MC/MD procedure. The absolute minimum of $E_{GB}$ is set to 0.

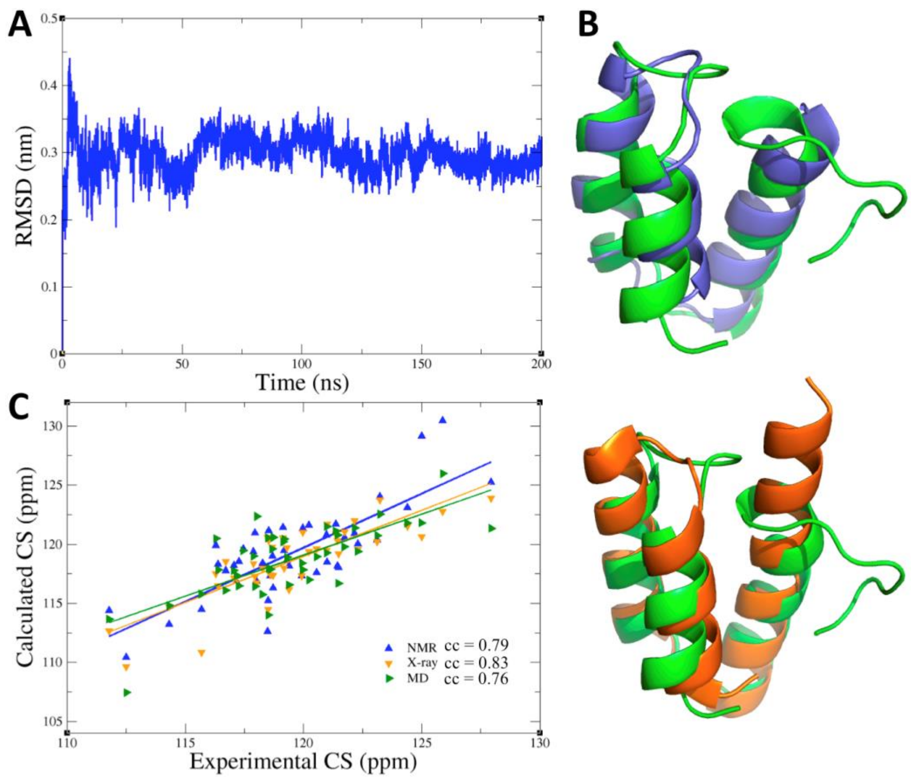


**Figure S7** MD simulations of $P_{XD}$ in water. **A** Backbone root-mean-square deviation (RMSD) relative to the initial structure of $P_{XD}$ plotted as a function of simulated time. **B** Superposition between the MD structure (green) and either the NMR structure (blue, PDB ID: 2K9D[1], top) or the X-ray structure (orange, PDB ID: 1OKS[24], bottom). **C** Calculated and experimental backbone N chemical shifts (CS)[1] of $P_{XD}$ as observed in the three structures in **B** The correlation coefficients (cc) are also reported. Color coding as the same in **B** Chemical shifts were calculated using the SHIFTX package[25].

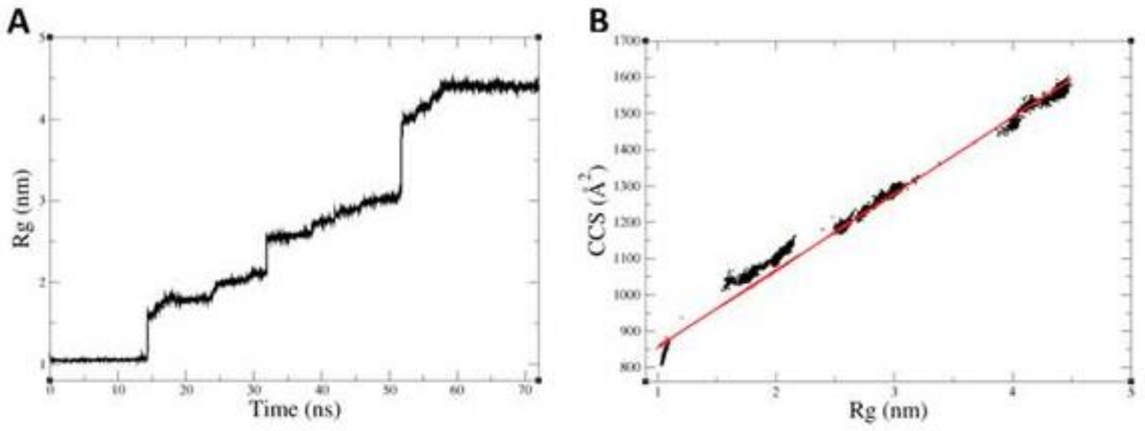


**Figure S8** SMD simulations of $[P_{XD}]^{5+}$. **A** Radius of gyration ($R_g$) plotted as a function of simulated time. **B** Correlation between $R_g$ and CCS. The correlation coefficient is 0.99.

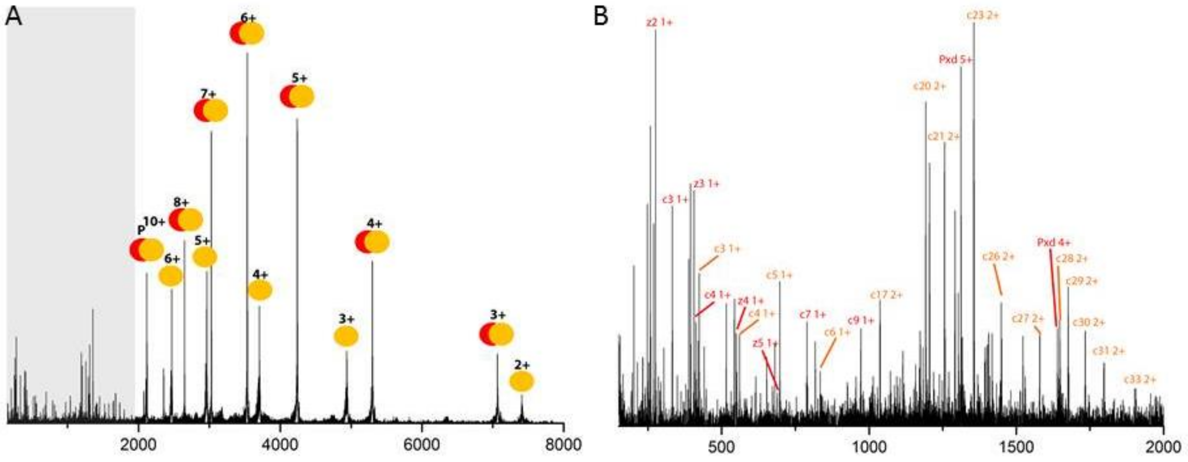

**Figure S9. A** Mass spectrum from top-down ETD experiment of the $10^+$ charge state of the $N_{TAIL}$-$P_{XD}$ complex. Pairs of orange and red circles represent the charge reduced precursor complex, whereas sole orange circles represent dissociated $N_{TAIL}$. ETD fragments were observed in the highlighted (grey) area. **B** ETD fragment spectrum with assignment of the most dominant fragments observed, along with dissociated $P_{XD}$.

**Table S1.** Information on the plateaus observed in the calculated CCS from the SMD simulation of $[P_{XD}]^{5+}$ in the gas phase.

| Plateau | Time period | Average CCS (Å$^2$) |
|---|---|---|
| 1 | 1 ns to 14 ns | 829±9 |
| 2 | 15 ns to 24 ns | 1055±11 |
| 3 | 25 ns to 30 ns | 1105±13 |
| 4 | 32 ns to 38 ns | 1189±9 |
| 5 | 42 ns to 50 ns | 1275±16 |
| 6 | 58 ns to 72 ns | 1563±10 |
